# *In silico* prediction of cellular organelles from computationally super-resolved (SR) phase-modulated optical micrographs

**DOI:** 10.1101/2025.09.14.675997

**Authors:** Shiraz S. Kaderuppan, Anurag Sharma, Muhammad Ramadan Saifuddin, Wong Wai Leong, Eugene, Wai Lok Woo

## Abstract

Differential interference contrast (DIC) & phase contrast microscopy (PCM) represent 2 widely-assimilated optical microscopical imaging modalities often employed for live cell imaging (LCI) applications. Although both of these approaches have marked advantages over traditional brightfield (BF) microscopy (they don’t require staining and provide real time visualization of cellular dynamics & metabolomics), there is a need to often supplement these modalities with epifluorescent microscopical imaging for the detection of specific organelles and structures *in vivo* (a process known as fluorescence combination/*colocalization microscopy*). Nonetheless, epifluorescent microscopy (both widefield and confocal) rely on the use of fluorophores (specialized fluorescent molecules and/or fluorescent proteins) for labelling key structures and features in the cell, which are prone to several issues, including photobleaching or cross-talk (amongst others). In this context, we seek to develop a novel deep neural network (DNN)-based approach aimed at predicting the location of 3 specific organelles (the cell nucleus, mitochondria & Golgi Apparatus) from acquired PCM & DIC images which have been super-resolved (SR) using our previously developed (& published) O-Net & Θ-Net model architectures. The model-generated images depict relatively close correlation with the ground truth images, implying that the O-Net & Θ-Net model architectures serve as viable frameworks for DNN model sculpting and training for both the purposes of image SR in optical microscopy as well as feature localization in these SR micrographs. We thus surmise on the potentiality of our proposed O-Net & Θ-Net DNN architectures for the development of models to be deployed in fully-automated image analysis pipelines in biomedical and healthcare diagnostics in the near future.

## Introduction

Zernike phase contrast microscopy (PCM) & differential interference contrast (DIC) microscopy have long been regarded as fundamental phase-modulated optical microscopical imaging modalities in live cell imaging & analysis. A key reason underlying this refers to the fact that both PCM & DIC microscopies are able to resolve miniscule features in cells [such as vesicles, mitochondria, the endoplasmic reticulum (ER) & Golgi Apparatus, amongst others] due to the relative differences in refractive indices (as compared to that of the surrounding cytosol). In PCM, light passing through these organelles becomes relatively retarded in phase (often by ¼ λ, or less) as compared to light waves transmitting through the surrounding cytosol, while these same waves undergo a further ¼ λ phase shift upon being transmitted through the phase plate in the objective [1]. When the optical path difference (OPD) between these waves approaches nλ or (n + ½) λ, they are able to undergo both constructive and destructive interference [1], highlighting the slight differences in refractive index between these said structures as stark contrast variations (see Figure 1 below for details). Significant changes encountered in the refractive index gradient are manifested as bright edges (in positive PCM) or dark rings (in negative PCM), giving rise to the characteristic halo observed around the feature of interest in PCM.

**Figure 1:**
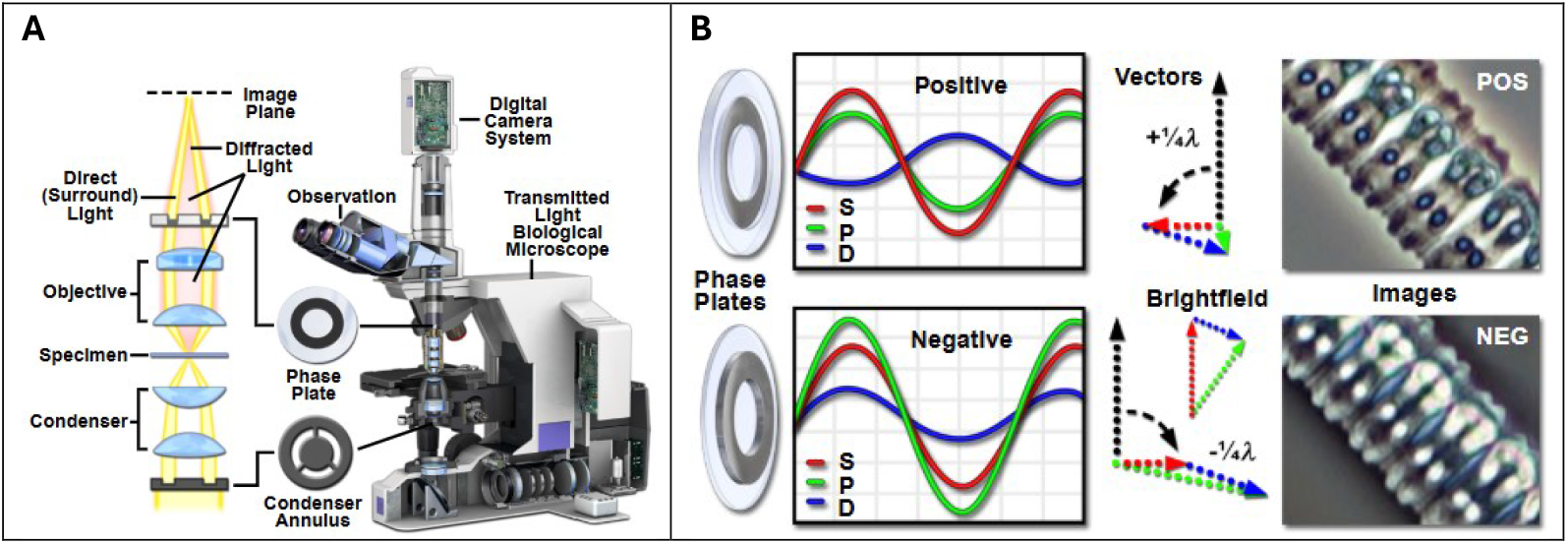
**A** Schematic showing the optical train in an upright phase contrast microscope. **B** An image of the green algae ***Zygnema*** acquired under both positive (top) & negative (bottom) PCM. Figure Source: [1].

DIC microscopy relies on a similar approach – to highlight slight differences in refractive index as pseudo-relief gradients. Light is transmitted through a birefringent Normaski/Wollaston prism in the condenser, where the O-& E-rays are split by a distance of ∼150-600nm in the specimen (object) plane [2]. The rays pass through 2 separate points in the sample (where one ray becomes retarded in phase relative to the other), before being recombined in another Wollaston prism after traversing the microscope objective. Recombination of the phase-shifted O-ray relative to the E-ray (or vice versa) leads to the visual perception of pseudo relief gradients at different points of the specimen, although it would be prudent to mention that these do not represent the *actual* topology of the specimen (see Figure 2 below for details).

**Figure 2:**
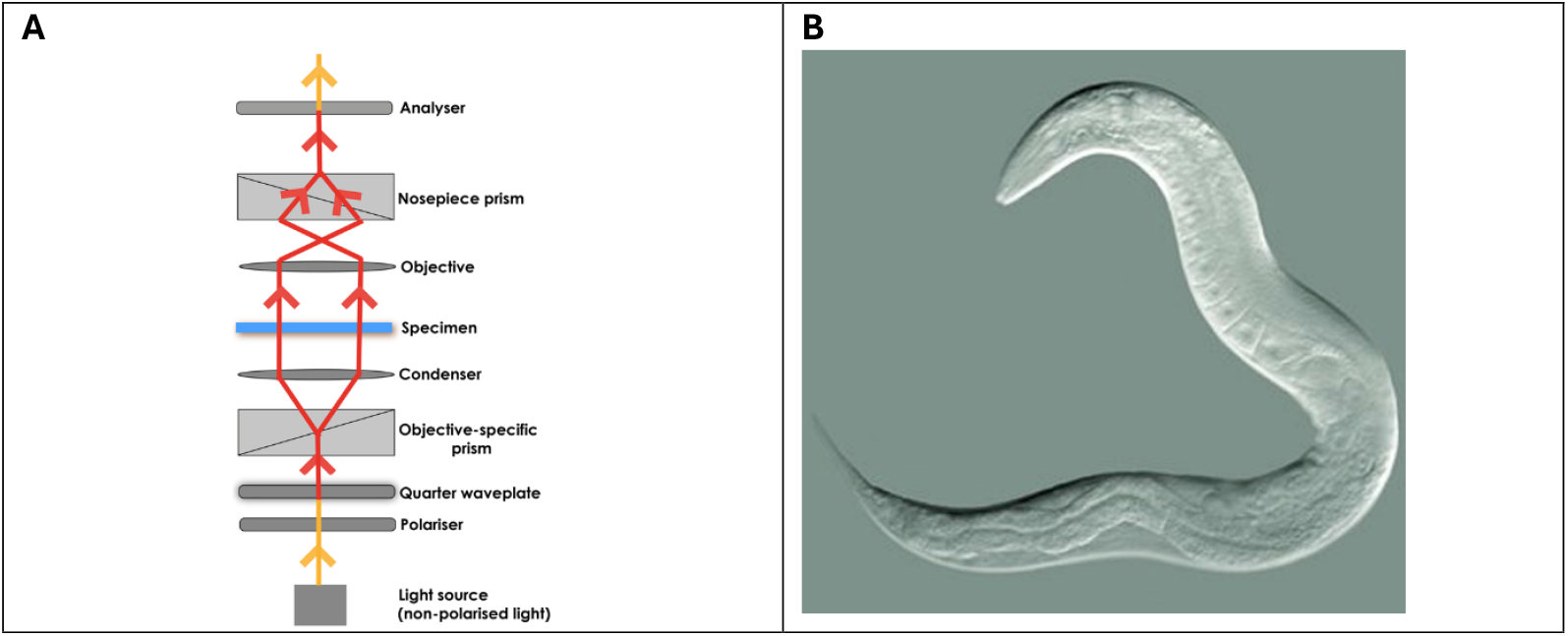
**A** Schematic showing the optical train of an upright DIC microscope. **B** An image of *C. elegans* acquired under DIC microscopy. Figure Source: [3].

Both PCM & DIC microscopy are highly sensitive microimaging modalities, being able to differentiate between features having an OPD < λ/100 [4]. Nonetheless, a characteristic setback faced with each of these techniques in LCI refers to their inability to distinguish between individual organelles in the cell. Due to this, research studies which employ either of these modalities would often have to utilize epifluorescent microscopical imaging in conjunction with these approaches for the identification of specific organelles or structures such as the plasma membrane, actin cytoskeleton or the nucleus) *in vivo* – an aspect known as *fluorescence combination/colocalization microscopy* [5]. In this regard, we sought to develop a purely computational approach employing deep neural networks adopted from our presently-developed O-Net [6] & Θ-Net [7] model architectures (as depicted in Figure 3 below) to facilitate the localization of 3 organelles of interest (namely the nucleus, mitochondria and Golgi Apparatus) from PCM or DIC micrographs. These micrographs are first super-resolved *in silico* through a different set of models (also adopting our previously-proposed O-Net network architecture) prior to being subjected to another set of models trained on the O-Net & Θ-Net frameworks. The Methodology section describes our approach in preparing the dataset for training the models as well as the model training methodology, while the Results section of this article depicts our findings gleaned from the said models.

**Figure 3:**
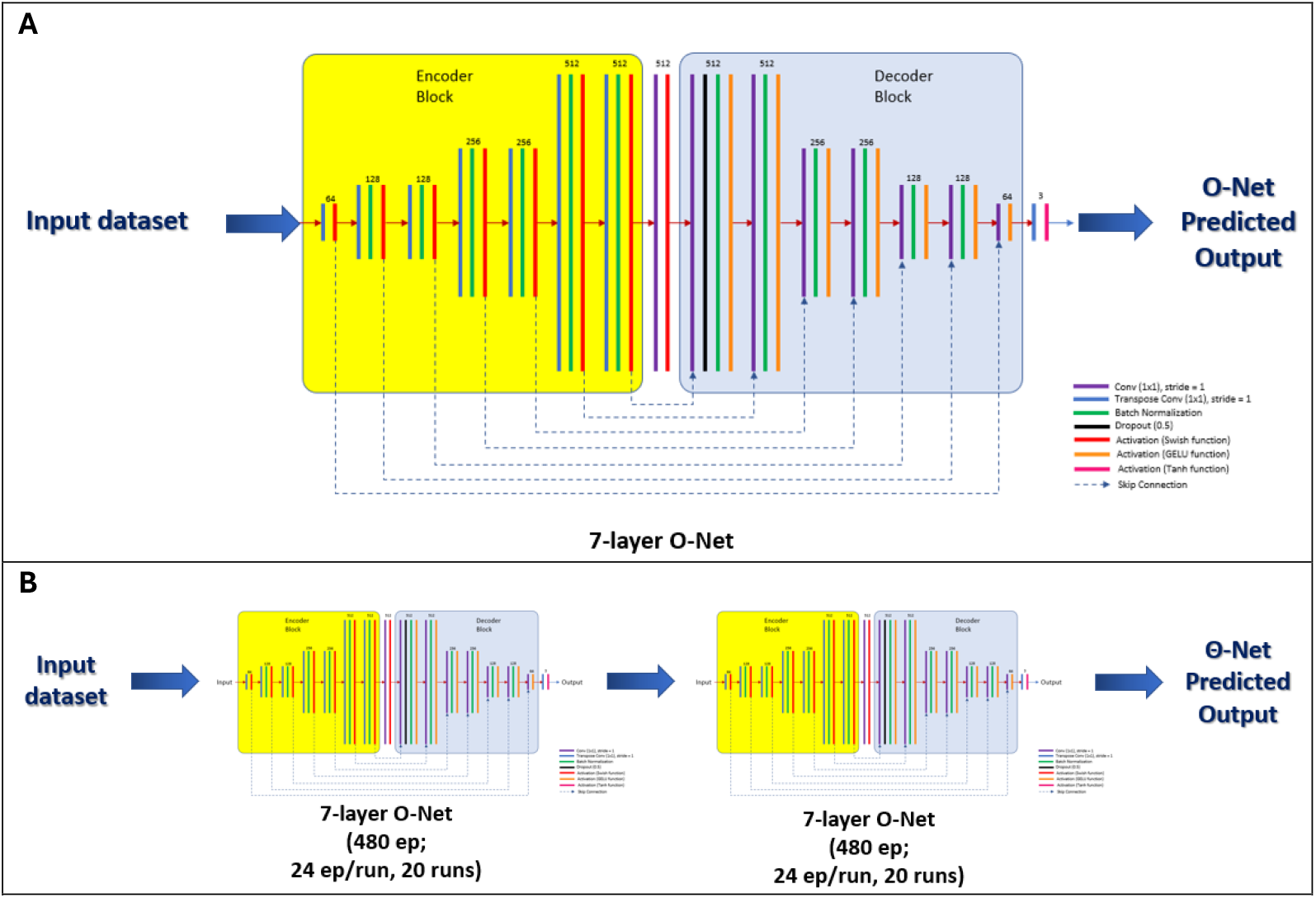
Schematics outlining the O-Net (**A**) & 2-node Θ-Net (**B**) network architectures as utilized in this present study. Further details pertaining to each of these DNN frameworks are described in our prior publications (see [6] & [7] for details).

**Figure 4:**
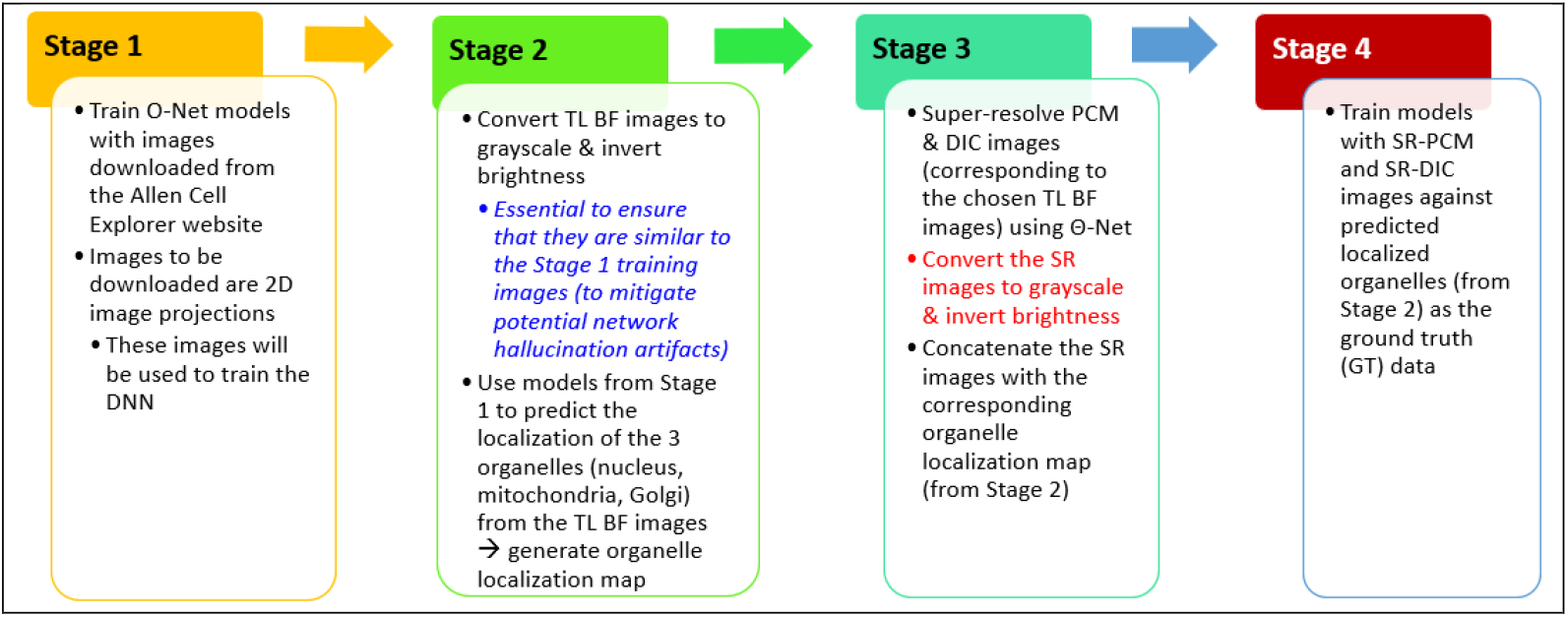
Pipeline showing the 4 stages as utilized in the current context of organelle detection & localization in the present study. The details pertaining to each of these individual stages are described in the following sections.

**Figure 5:**
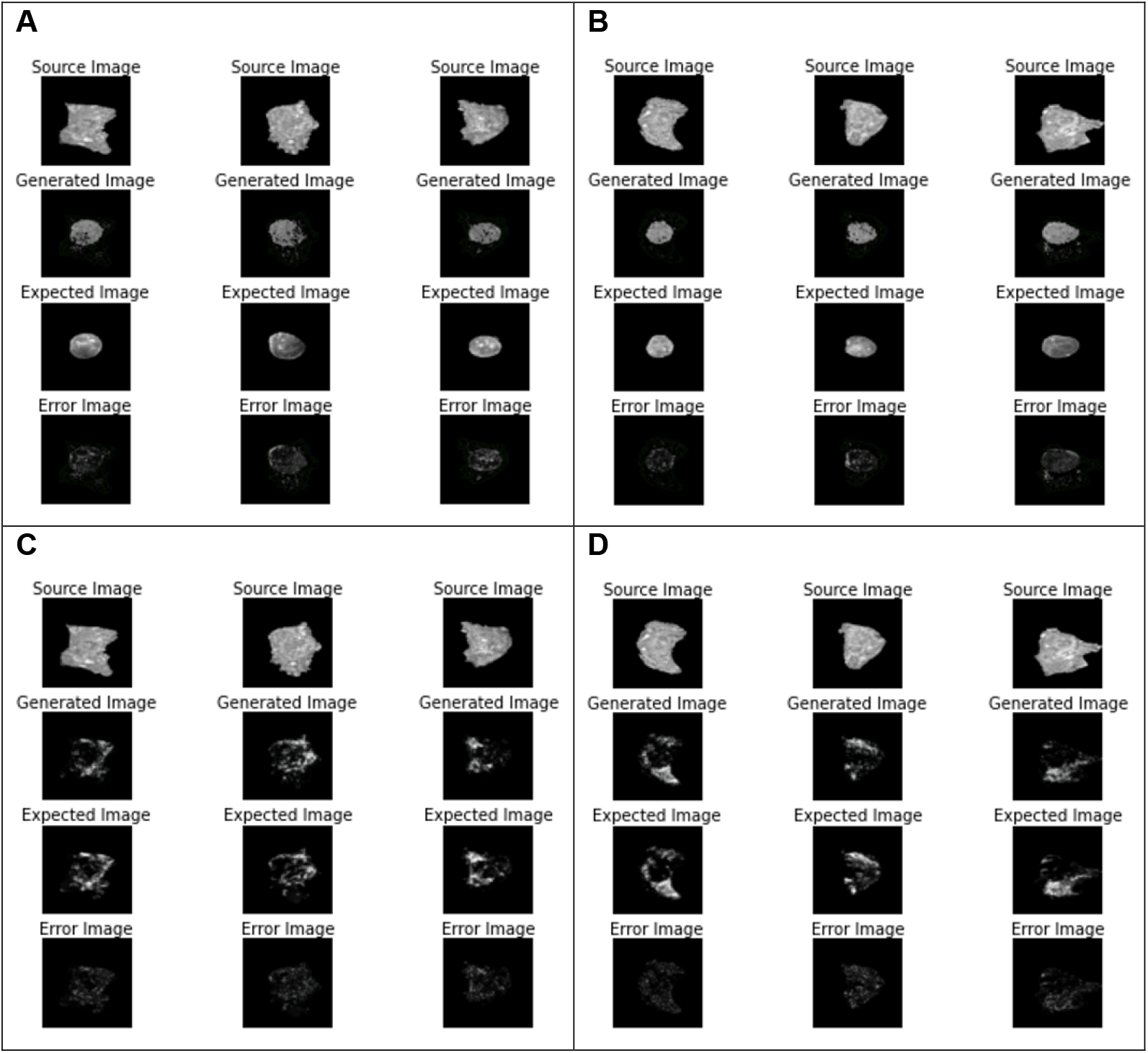
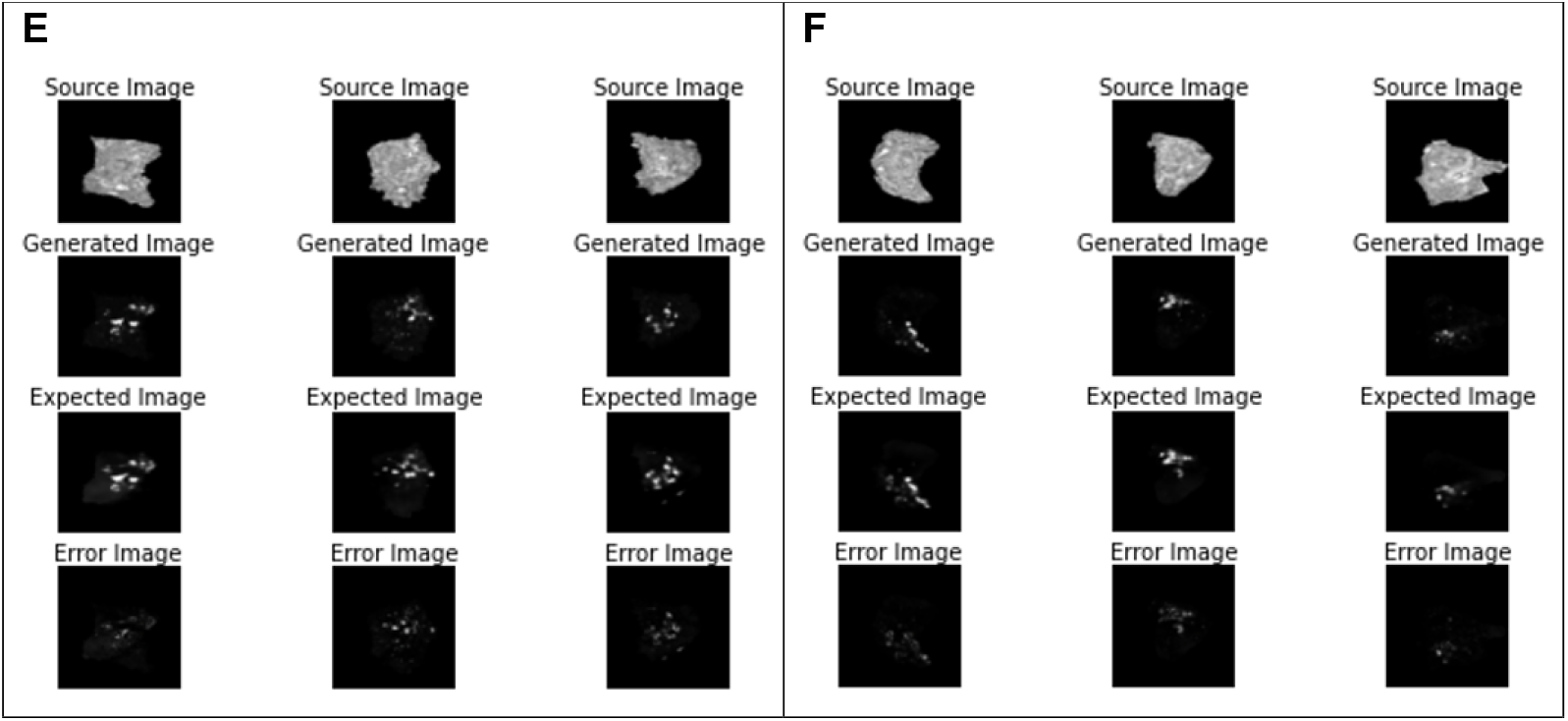
Validating the Stage 1 model performance using images obtained from the training dataset (panels **A, C & E**) as well as the untrained (validation) image dataset (panels **B, D & F**) for the Nucleus (panels **A & B**), Mitochondria (**C & D**) and the Golgi Apparatus (**E & F**) respectively. All of the 3 models assayed here were based on a single 7-layer O-Net architecture, with the Nucleus, Mitochondria and Golgi Apparatus models being trained over 4,114,800, 4,011,930 & 3,909,060 iterations respectively. For each panel, the model output (**Generated Image**) is compared against the **Expected** (ground truth) image, where the **Error Image** describes the absolute difference in pixel intensities between the **Generated** and **Expected** (ground truth) images.

## Methodology

Training of the DNN models as described in this study proceeded in 4 Stages as depicted in the flow chart below:

### Stage 1

We prepared our first round of model training with the image dataset of these 3 organelles (the nucleus, mitochondria & Golgi Apparatus) as downloaded from the Allen Cell Explorer website ([8] & [9]). Using these images (which consisted of the transmitted light brightfield images of the individual cells coupled with the 2D-projected epifluorescent images of the targeted organelles), we trained our models adopting the O-Net architecture. For the Golgi Apparatus & mitochondria datasets, the models were trained over 200 epochs (20 runs of 10 ep/run), where each epoch consisted of 20,574 iterations (hyperparameter switching was performed after the 12^th^ run). For the Nucleus dataset, we utilized *transfer learning*, adopting a training seed of a model previously trained with a quad-image dataset of the downloaded nuclei images.

### Stage 2

Diascopic brightfield (BF), PCM & DIC images of selected regions of interest (RoIs) are acquired as described in [6]. The BF images are converted to their 8-bit grayscale equivalents & their brightness inverted within ImageJ (to make these images similar to that used for training the models in Stage 1 previously). These images were the fed into the models developed during Stage 1 above to generate the corresponding putative fluorescent images for the cellular nucleus, mitochondria & Golgi Apparatus (i.e. a predicted organelle localization map).

### Stage 3

The PCM & DIC images of the RoIs as described in Stage 2 above are super-resolved using the O-Net models as published previously in [6]. The SR images obtained are then converted to 8-bit grayscale and concatenated with their corresponding organelle localization maps (generated from Stage 2), resulting in a total of **6 datasets** (SR PCM-Nucleus, SR DIC-Nucleus, SR PCM-Mitochondria, SR DIC-Mitochondria, SR PCM-Golgi Apparatus & SR DIC-Golgi Apparatus).

### Stage 4

The datasets from Stage 3 are then used to train a separate set of O-Net & Θ-Net models. For the SR DIC-Nucleus & SR DIC-Mitochondria images, we developed our models using a 7-layer O-Net architecture (20 runs of 24 epochs/run, with hyperparameter switching initiated after the 12^th^ run as in Stage 1 previously). For the other 4 datasets (SR PCM-Nucleus, SR PCM-Mitochondria, SR PCM-Golgi Apparatus & SR DIC-Golgi Apparatus), a 2-node (7-layer) Θ-Net model architecture was employed. Nonetheless, at the current juncture, results obtained from the Θ-Net models are only presented for the SR PCM-Golgi Apparatus & SR DIC-Golgi Apparatus datasets; for the SR PCM-Nucleus & SR PCM-Mitochondria datasets, the O-Net model results are shown in this study.

## Results

The results obtained through the use of the afore-mentioned models are depicted as follows:

### Stage 1

#### Stage 4

The following images describe the model output & its comparison against the **Expected** (ground truth) images, where the blue-bounded RoIs indicate features having a reduced SNR, while the green RoIs indicate supposedly unresolved features.

## Discussion

From the results obtained for Stage 1 above, the models exhibit a relatively high viability with the **Generated** images closely resembling the **Expected** (ground truth) images. This suggests a high potentiality for these models to be used in the subsequent stage (Stage 4) for generating the ground truth images to train a separate set of models for localizing the organelles *in vivo*.

For Stage 4, the model-generated images demonstrate a relatively high similarity to the **Expected** (ground truth) images for the Golgi Apparatus (Figure 8) where a 2-node Θ-Net architecture is used. This is particularly true for the 2^nd^ node **Gen** images, which amplify image SNR relative to the 1^st^ node **Gen** images. Fundamentally, features which are clearly identifiable within the **Source** and the **Expected** images are also evident in the model-generated images, although there do still exist some features which are not evident in the **Source** image but are indicated in the **Expected** image (these are highlighted in the green-bounded RoIs within these images). Closer inspection of these discrepancies reveal that this might actually be an artifact arising from improper elucidation of the Golgi Apparatus from the BF images, suggesting that the stage 4 models are actually resilient to network overfitting and potential hallucination artifacts that might arise during the model validation process.

For the predicted Mitochondria distribution in Stage 4 (Figure 7), a similar trend is noted where the intensity of the pixels putatively indicating the mitochondria localization *in vivo* is generally reduced as opposed to the **Expected** images, with some potential mitochondria in these images not detectable within the **Generated** images. Again, comparing these images to the input **Source** image may suggest that the model is relatively resilient to over-fitting, although there is room for further improvement of the model performance during training as well. The same observations are replicated in the Nucleus dataset as well (Figure 6), with the model performance for the PCM dataset being generally lower than that of the model performance for the DIC dataset (for the untrained/validation images), suggesting further room for improvement of the model training process.

**Figure 6:**
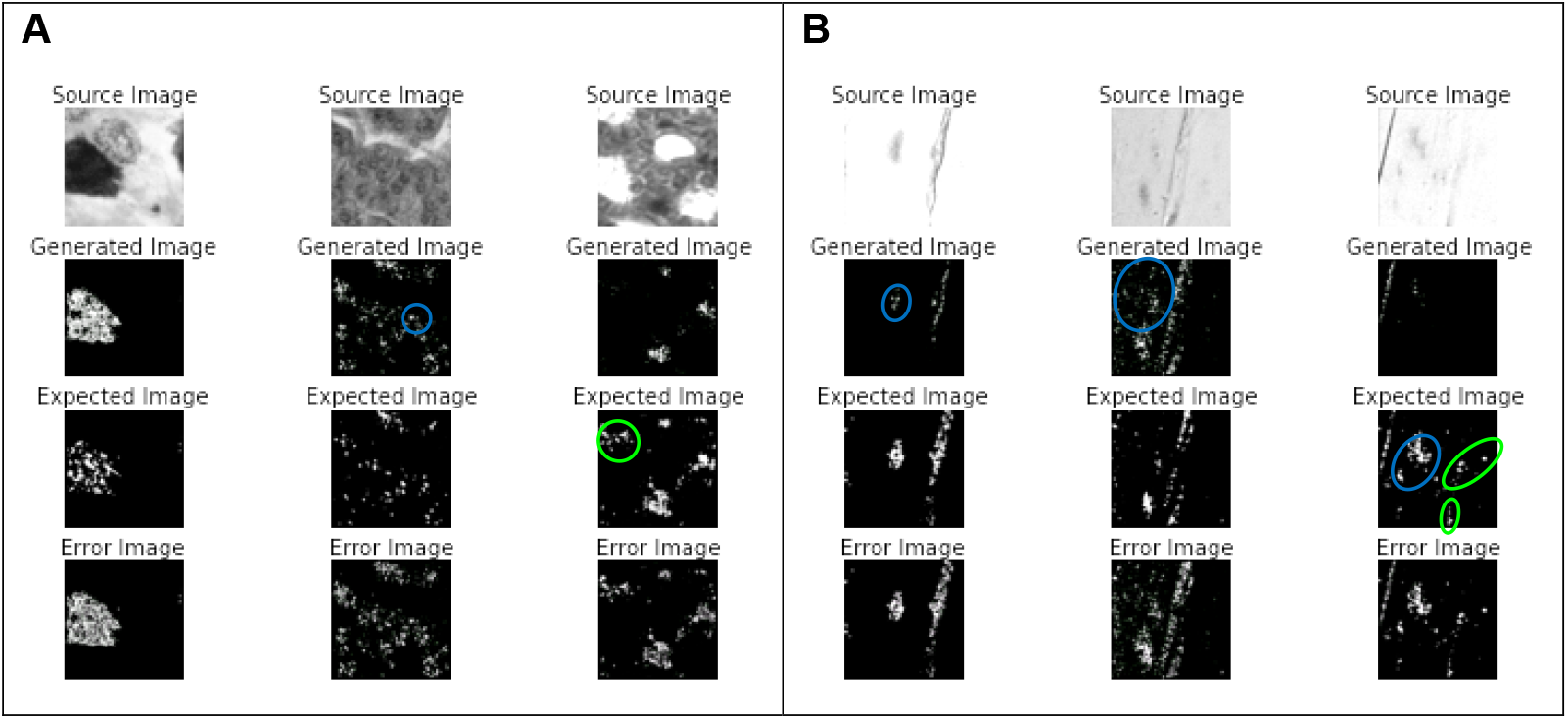
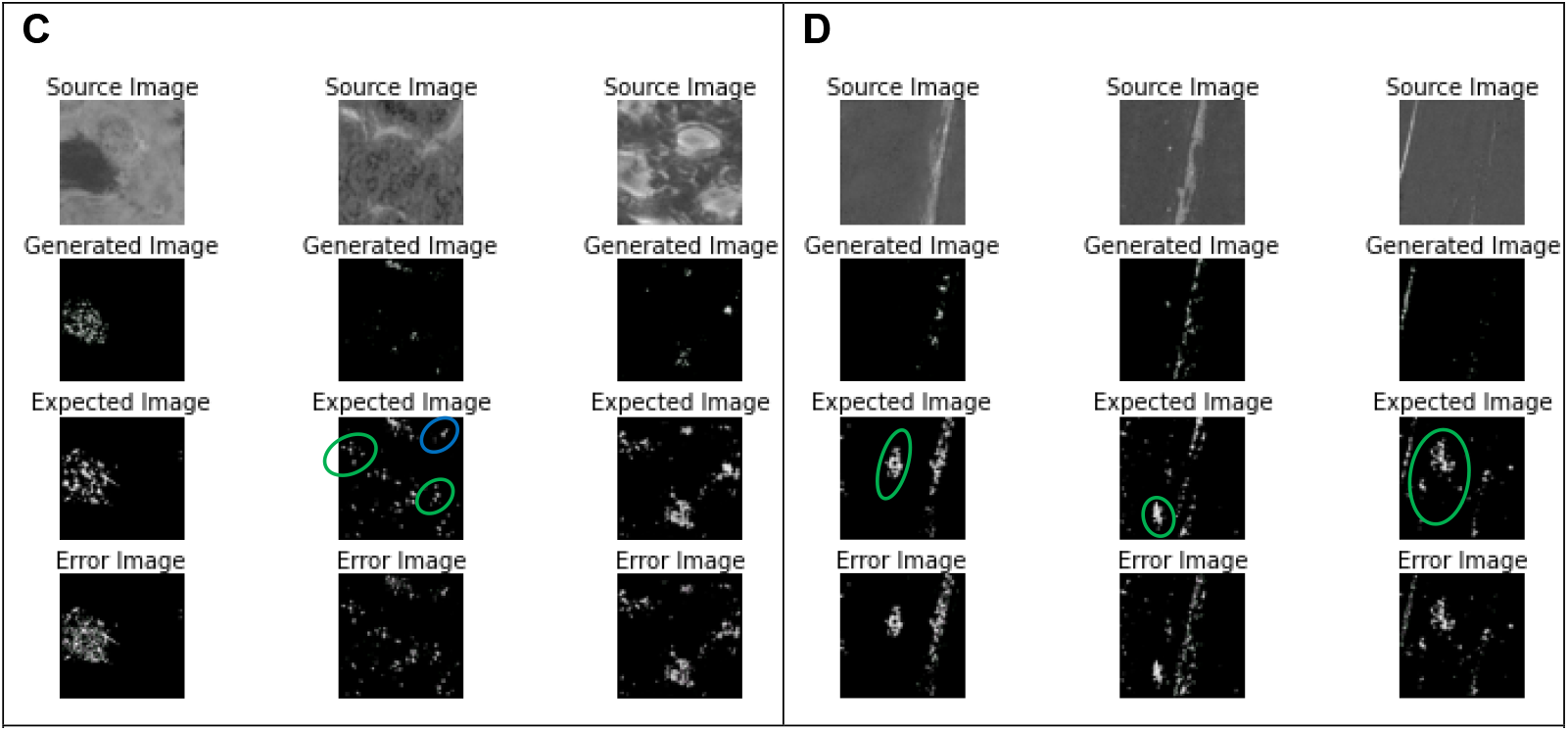
Validating the Stage 4 model performance using images obtained from the training dataset (panels **A & C**) as well as the untrained (validation) image dataset (panels **B & D**) for the SR DIC-Nucleus (panels **A & B**) and the SR PCM-Nucleus (**C & D**) datasets respectively. Here, the models utilized were founded on a single 7-layer O-Net architecture, trained over 1,187,280 & 419,040 iterations for the SR DIC-Nucleus & SR PCM-Nucleus datasets respectively. Also (as with Stage 1 previously), in each panel, the model output (**Generated Image**) is compared against the **Expected** (ground truth) image, where the **Error Image** describes the absolute difference in pixel intensities between the **Generated** and **Expected** (ground truth) images.

**Figure 7:**
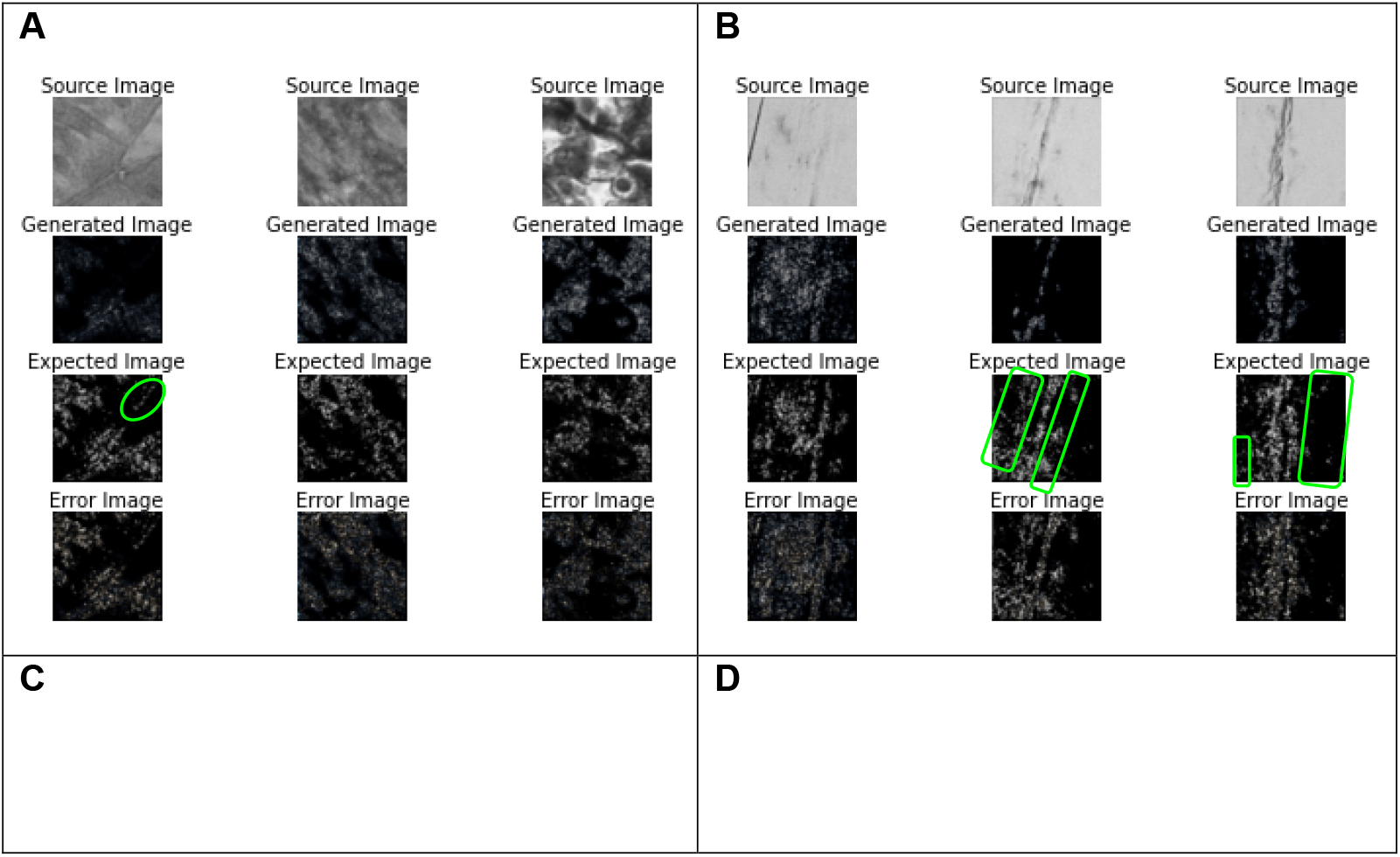
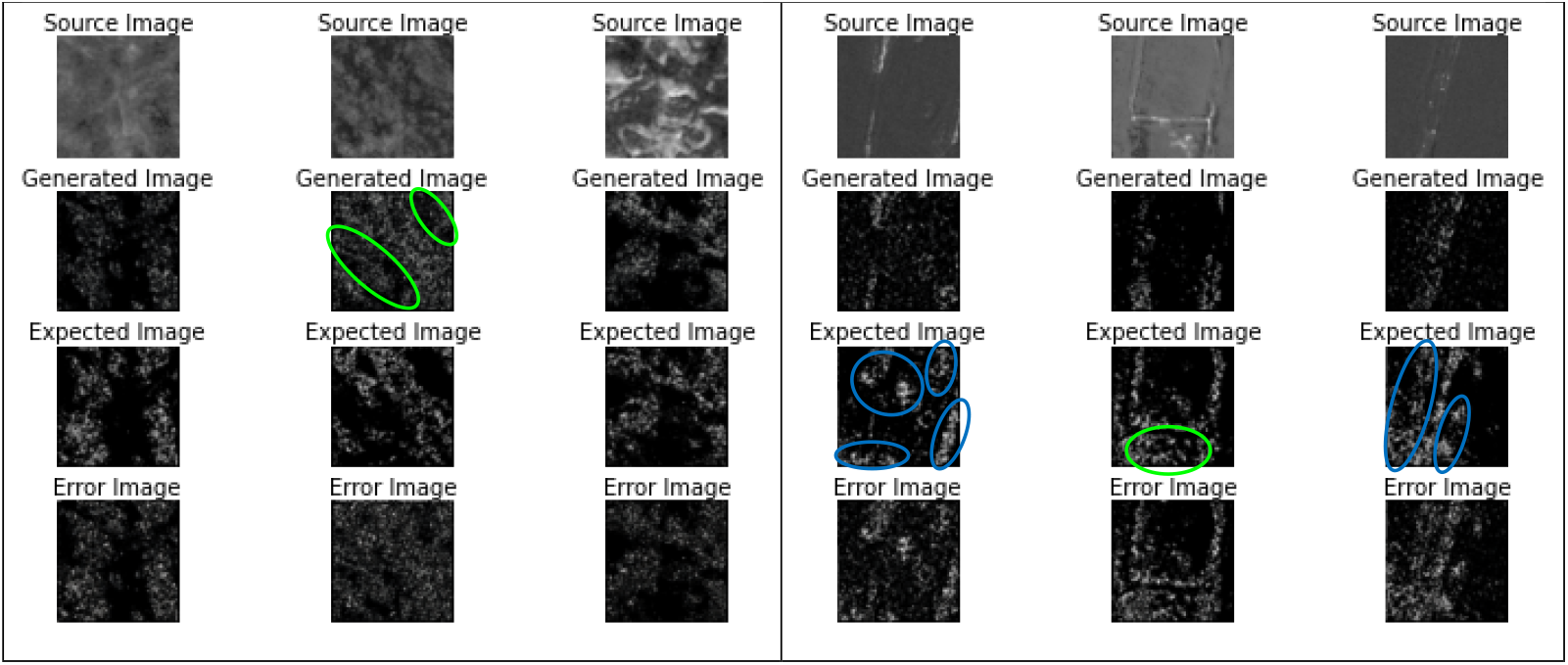
Validating the Stage 4 model performance using images obtained from the training dataset (panels **A & C**) as well as the untrained (validation) image dataset (panels **B & D**) for the SR DIC-Mitochondria (panels **A & B**) and the SR PCM-Mitochondria (**C & D**) datasets respectively. Here too, the models adopted a single 7-layer O-Net architecture, trained over 1,373,520 & 1,338,600 iterations for the SR DIC-Mitochondria & SR PCM-Mitochondria datasets respectively. As was done previously, the differences between the model output (**Generated**) against the **Expected** (ground truth) images are described in the **Error Image**, which depicts the absolute difference in pixel intensities between these 2 images.

**Figure 8:**
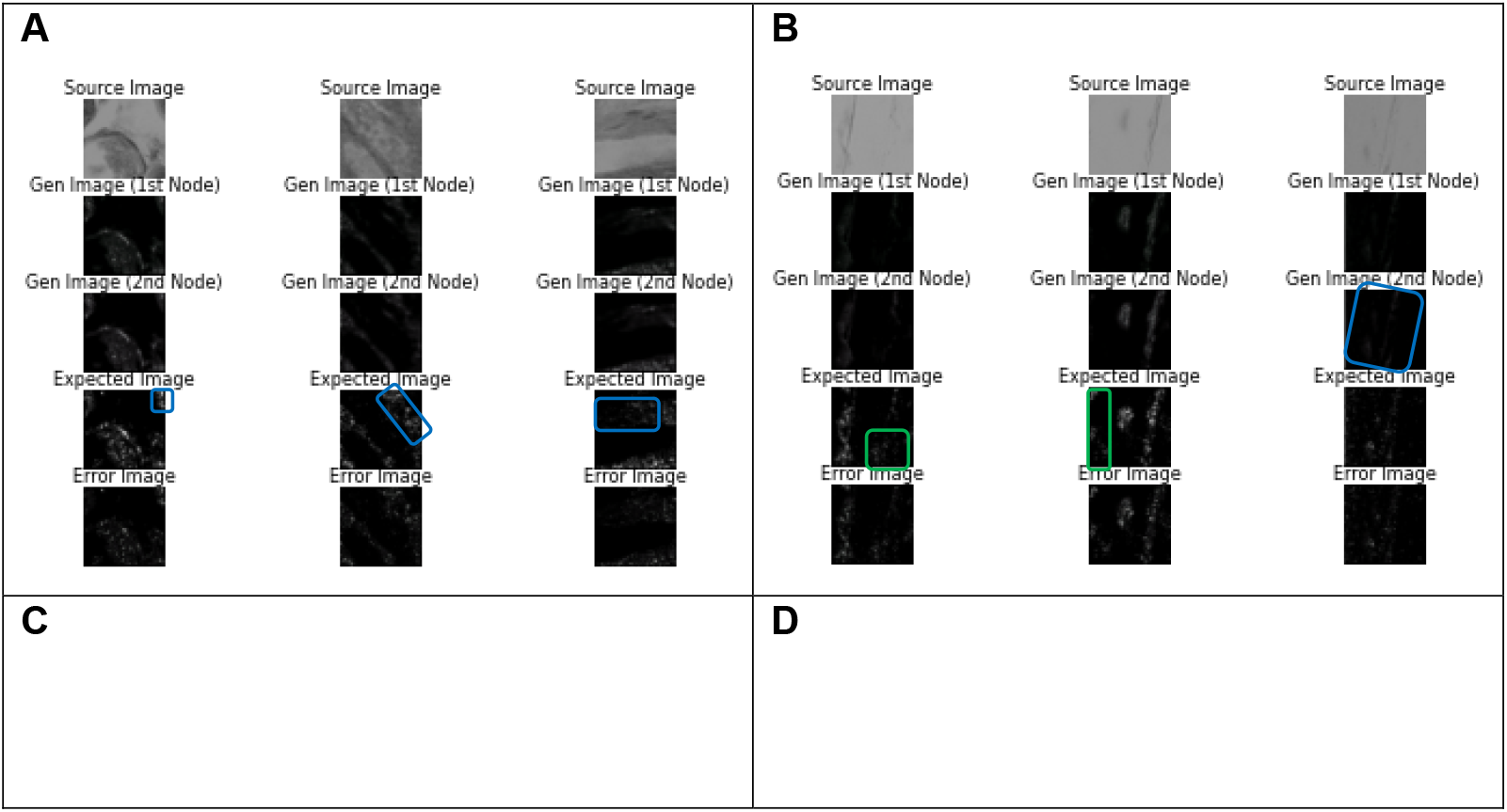
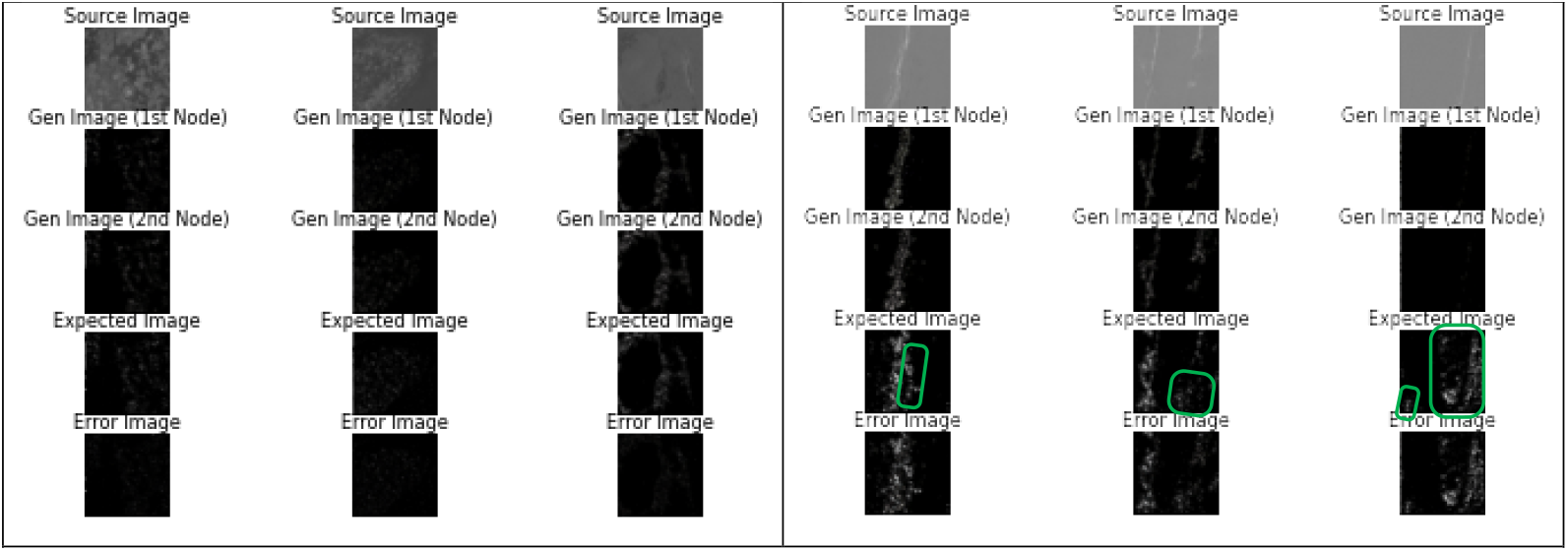
Validating the Stage 4 model performance using images obtained from the training dataset (panels **A & C**) as well as the untrained (validation) image dataset (panels **B & D**) for the SR DIC-Golgi Apparatus (panels **A & B**) and the SR PCM-Golgi Apparatus (**C & D**) datasets respectively. Here, the models utilized were founded on a 2-node Θ-Net architecture (each node comprising of a 7-layer O-Net), trained over 523,800 & 1,280,400 iterations (1^st^ node) and 1,233,840 & 1,059,240 iterations (2^nd^ node) for the SR DIC-Golgi Apparatus & SR PCM-Golgi Apparatus datasets respectively. Also (as with Stage 1 previously), in each panel, the model output (**Generated Image**) is compared against the **Expected** (ground truth) image, where the **Error Image** describes the absolute difference in pixel intensities between the **Generated** and **Expected** (ground truth) images.

### Current limitations & Potential areas for future exploration & advancement

The presently-proposed study describes a novel approach to localizing some key cellular organelles of significant research interest (namely the nucleus, mitochondria & Golgi Apparatus) in a couple of non-fluorescent phase-modulated microimaging modalities (i.e. DIC & PCM). The results gleaned from our models demonstrate the feasibility of both O-Net & Θ-Net as viable DNN architectures for model training in the space of label-free in silico organelle detection and localization. Nonetheless, it would also be prudent to emphasize at this juncture that the models adopted in our current study do indeed have a few potential areas for further explorations and improvement, namely the use of a fluorescent dataset together with the PCM & DIC images (rather than a BF image dataset). This would also alleviate the number of steps in the current workflow, thereby optimizing the process while reducing potential errors that may arise. Another area for further exploration would be to increase the types of organelles and structures being localized *in vivo*, such as the actin cytoskeleton, rough/smooth endoplasmic reticulum (r/sER), lysosomes, etc.

## Conclusion

The potential of our proposed O-Net & Θ-Net model architectures have been demonstrated in this study to be capable of localizing organelles of interest within a series of non-fluorescent micrographs. The accuracy of the model though are still subject to further refinements, with improvements in the dataset acquisition coupled with a reduction in the number of Stages required in the model training workflow. Application-wise, these models may be postulated to have far-reaching implications across industry, including (amongst others) drug perturbation studies as well as cellular metabolomics, just to name a few. In this regard, the *in silico* identification & localization of cellular organelles and even proteins of interest remain potential avenues for future research and exploration.

